# Cryo-EM visualization of an active high open probability CFTR ion channel

**DOI:** 10.1101/274316

**Authors:** Jonathan F. Fay, Luba A. Aleksandrov, Timothy J. Jensen, Liying L. Cui, Joseph N. Kousouros, Lihua He, Andrei A. Aleksandrov, Drew S. Gingerich, John R. Riordan, James Z. Chen

**Affiliations:** University of North Carolina, Chapel Hill; Oregon Health & Science University

## Abstract

The Cystic fibrosis transmembrane conductance regulator (CFTR) anion channel, crucial to epithelial salt and water homeostasis, and defective due to mutations in its gene in patients with cystic fibrosis is a unique member of the large family of ATP-binding cassette transport proteins. Regulation of CFTR channel activity is stringently controlled by phosphorylation and nucleotide binding. Structural changes that underlie transitions between active and inactive functional states are not yet fully understood. Indeed the first 3D structures of dephosphorylated, ATP-free and phosphorylated ATP-bound states were only recently reported. Here we have determined the structure of inactive and active states of a thermally stabilized CFTR with very high channel open probability, confirmed after reconstitution into proteoliposomes. The unique repositioning of the TMHs and R domain density that we observe provide insights into the structural transition between active and inactive functional states of CFTR.

**Highlights:**

- Structures of thermostabilized avian CFTR in dephosphorylated or phosphorylated forms at 4.3 Å and 6.6 Å resolution, respectively.
- Conformational differences of transmembrane helices 7 & 8 compared to zebra fish and human CFTR structures reveal an extracellular vestibule that may provide anion access to the pore.
- R-domain density appears to “plug” the intercellular vestibule in the dephosphorylated avian CFTR cryo-EM map.

## Introduction

Cystic fibrosis (CF) is a heritable disorder that leads to dysfunction of the respiratory, digestive and reproductive systems. This systemic dysfunction is caused by mutations in the CF transmembrane conductance regulator (CFTR) gene, which compromise CFTR protein folding and function. Although, CF pharmacotherapies have advanced over the past decade, patients diagnosed with CF only have a median life expectancy of around 40 years. All of the somewhat effective small molecule CFTR modulators found to date have been discovered in cell based functional screens and their interactions with the protein have not yet been fully characterized (Ren, Grove et al. 2013). Attempts at structure based design to improve such compounds or discover new ones has so far been restricted to CFTR homology models generated using distantly related ABC exporter templates (Dalton, Kalid et al. 2012; Rahman, Cui et al. 2013; Mornon, Hoffmann et al. 2015; Das, Aleksandrov et al. 2017). Until very recently (Zhang and Chen 2016; Liu, Zhang et al. 2017; Zhang, Liu et al. 2017) no experimentally determined structural information was available because of difficulties in obtaining homogeneous and stable purified protein.

Studies initially of the most common cystic fibrosis causing mutation, the deletion of a single amino acid (phe508) from the first nucleotide binding domain (NBD1) had revealed not only that the primary impact of the mutation was to severely impair conformational maturation, lifetime and thermodynamic stability (TS) of the full length protein but also more recently that the wild-type protein had marginal stability (Aleksandrov, Jensen et al. 2015).

Previously, we rationally increased the stability of wild-type human CFTR by the modification of mobile segments of NBD1 and substitution of a catalytic residue In NBD2 involved in ATP hydrolysis and found that the quantity and quality of the protein expressed and purified from mammalian cells was greatly improved (Aleksandrov, Jensen et al. 2015). Here we have expressed and purified a similarly modified chicken CFTR with still greater TS in order to resolve the conformation of both the phosphorylated active and dephosphorylated inactive CFTR channel by cryo-electron microscopy.

## Results

### Chicken CFTR expression, purification and reconstitution

We selected chicken CFTR (chCFTR) for structural studies because it is innately more thermally stable than the human channel protein (Aleksandrov, Kota et al. 2012) and the sequence similarity of chicken CFTR compared to human is ∼ 80% (notably higher than that of zebrafish ∼55%). Although combined stabilizing modifications of human CFTR increased it’s Tm from ∼44 °C to ∼53 °C, similar modifications of the inherently more stable chicken CFTR resulted in a Tm of ∼58 °C (Fig S1). The thermostabilized chicken CFTR (chTS) was stably expressed in baby hamster kidney (BHK) cells, adapted to suspension culture for large scale production as described (Aleksandrov, Jensen et al. 2015). As seen in Figure 1A microsomal membranes derived from chTS (ΔRI/1404S/1441X; illustrated in Fig S2B,) are devoid of CFTR channel function when treated with protein phosphatase, and channel activation occurs after treatment with protein kinase A (PKA).

**Figure 1.**
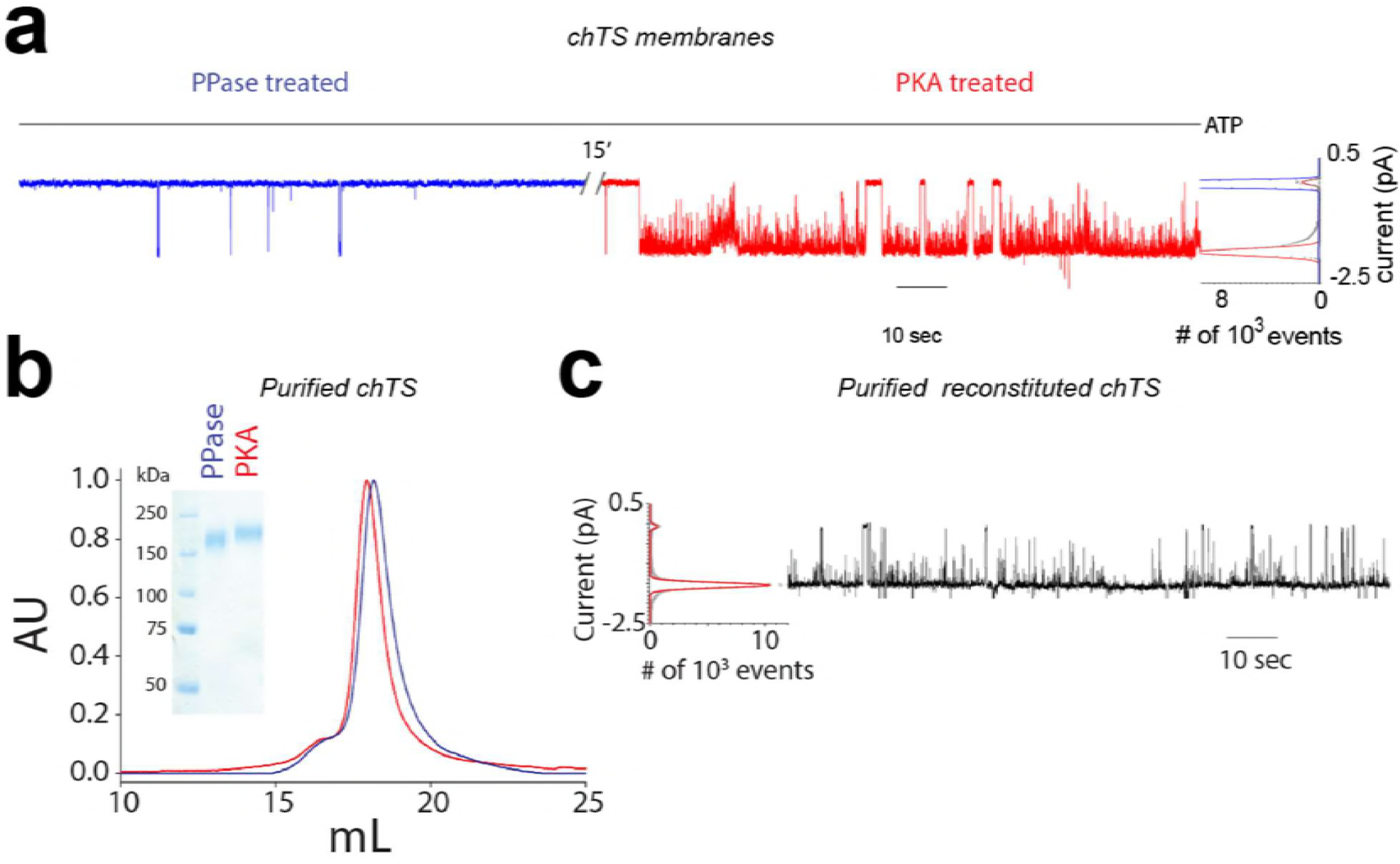
Thermostabilized Chicken CFTR is activated to a high open probability state by phosphorylation and this behavior is retained after purification. **a)** Single channel recordings of Chicken ΔRI/1404S/1441X/His construct (chTS) expressed in BHK microsomal membranes show the channel is inactive when dephosphorylated (PPase treated 1, blue) and nearly locked-open (Po=0.95) when phosphorylated (PKA treated 2, red). The open state probability calculated from histograms to the right are indicated for PPase treated (blue) and PKA treated (red) states. **b)** Fluorescent Size Exclusion Profiles of chTS for PPase or PKA treated CFTR (Blue and Red). Mg-ATP was present at 5 mM in the running buffer. Note the decrease in migration time for the phosphorylated species. inset SDS-PAGE analysis of IMAC purified dephosphorylated (PPase, blue) and PKA treated chTS (red). c) Single channel recording of PKA treated purified chTS reconstituted into proteoliposomes, resulting in a high open probability state similar to that in microsomes.

Microsomes were then solubilized in detergent and purified using ion metal affinity chromatography to produce pure protein in either a dephosphorylated or phosphorylated form. Biochemical analysis indicates the PKA treated purified chTS is phosphorylated, as evidenced by an electrophoretic gel shift, size-exclusion chromatography (SEC) shift (Fig 1B) and further confirmed by mass-spectroscopy (Fig S2A). Major PKA phosphorylation sites in the R-domain controlling channel activity were unphosphorylated after PPase treatment and phosphorylated after PKA exposure (Fig S2).

The purified chTS was then reconstituted into proteoliposomes and single channel recordings were performed. The conductance and high Po were nearly identical to those of the channel in the native microsomes from which it was purified (Fig. 1C) indicating that channel function is retained by the purified protein.

Moreover, these functional results with chTS are similar to those of hCFTR containing the corresponding catalytic histidine mutation in NBD2 (1402S) (Aleksandrov, Jensen et al. 2015) that has recently been confirmed to inhibit ATP hydrolysis (Yang, Hildebrandt et al. 2018). Other hydrolysis inhibiting mutations in NBD2 also cause high open probability channel activity (Vergani, Nairn et al. 2003).

### Structure of dephosphorylated chCFTR

Detergent purified dephosphorylated chTS in the presence of 5mM Mg-ATP eluted directly from metal affinity columns, was used for cryo-EM structure determination after concentration to 7 mg/ml, exchanged into buffer made up in D_2_O rather than H_2_O, and removal of glycerol and imidazole. From 753 micrographs (movie frames by K2, see the Method Section), 221,536 particles were selected, 165,541 particles were retained after “2D filtering” in RELION2 (Scheres 2012). Single-particle three-dimensional analysis of these remaining particles in cryoSPARC (Punjani, Rubinstein et al. 2017) revealed a single class with an overall resolution of ∼8 Å (FSC = 0.143). After multiple additional rounds of 3D classification, 30,219 particles could be reconstructed to a 3D map for which the FSC measurement at 0.143 indicated resolution to 4.3 Å for the masked protein (Fig 2A, Fig S3). For this reconstruction the transmembrane helices (TMH) and the connectivity between the helices are well resolved, with the exception of extracellular loop 4 (EL4).

**Figure 2.**
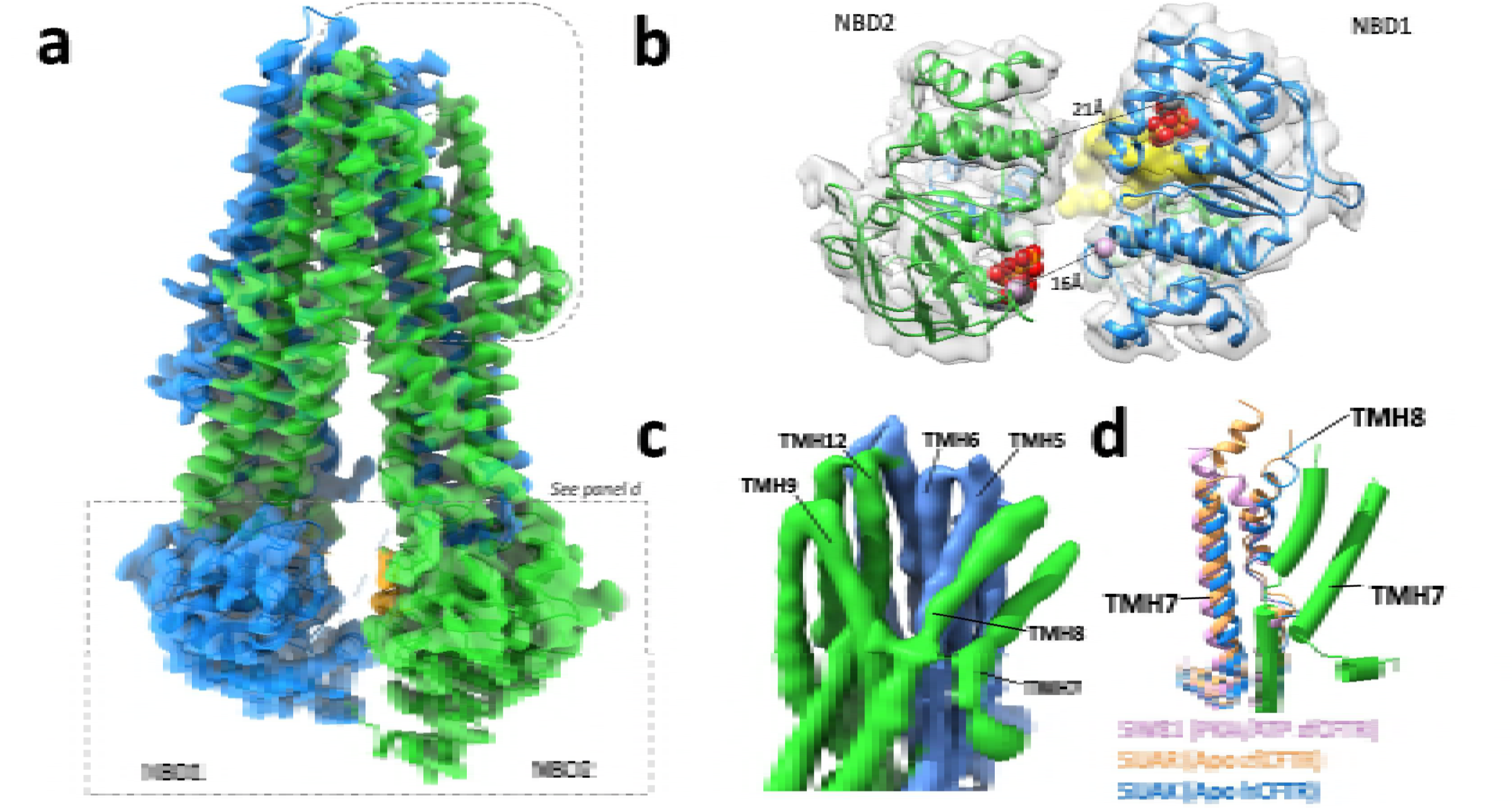
Electron density map of the dephosphorylated (PPase treated) and inactive chicken CFTR. **a)** Side view of the post processed cryo-EM map reconstruction obtained from three-dimensional refinement. Colored with MSD1/NBD1 as blue and MSD2/NBD2 as green. Note, the NBDs appear to not be in within ATP bridging distance (box). An extracellular vestibule is also seen, formed due to conformational transitions of TMH7 & 8 (rounded box). **b)** The distance between NBD1 Walker A residue G461 and NBD2 signature sequence residue S1347 is ∼21 Å (gray) compared to the 27 Å observed for zfCFTR (Zhang and Chen 2016). The NBD1 signature sequence S549 is separated from NBD2 Walker A G1247 by ∼16 Å (plum), approximately 2 Å closer than observed for corresponding residues of zfCFTR. High threshold R-domain density is shown in yellow. **c)** Shown is a side view of the unfiltered binned cryo-EM map obtained from three-dimensional reconstruction. An apparent extracellular vestibule is formed by TMH 5,6,9 & 12. **d)** Comparison of TMH7 & 8 positioning in chCFTR with that in PKA treated ATP bound zfCFTR (plumb, PDB 5W81), ATP free, dephosphorylated zf (sandybrown, PDB 5UAR) and human CFTR (blue, PDB 5UAK).

### Overall Architecture and low resolution features

Our map reconstruction (Fig 2) is similar to that of other published CFTR cryo-EM reconstructions (Zhang and Chen 2016; Liu, Zhang et al. 2017). However, some differences are apparent. Interestingly, although EL4 is not visible, at low threshold density, an extra density appears around the presumed EL4 location that could reflect the large complex N-linked oligosaccharide chains present in our protein(Aleksandrov, Kota et al. 2012) but not in those of Chen and colleagues, that contain only short core chains. At low resolution there is also additional density on the outside of NBD2 that would correspond to the amino acids that extend from TMH12 to NBD2. Also, we observe only the last helix portion of the so called “lasso motif” (Zhang and Chen 2016) while density for the first ∼40 residues of the N-terminus was less well resolved.

### Nucleotide Binding Domains (NBDs) are not within ATP bridging distance

Perhaps most notably, even in the RELION 2D class averages (Fig 3A) it is apparent that the NBD separation is intermediate between what has been observed in the dephosphorylated (Zhang and Chen 2016; Liu, Zhang et al. 2017) and the more recent ATP bound phosphorylated zebrafish CFTR cryo-EM models (Zhang, Liu et al. 2017). In chCFTR the NBD separation is not within ATP bridging distance. When aligned to NBD1 and membrane spanning domain 1 (MSD1) of the ATP free dephosphorylated zebrafish CFTR structure we see a rotation of the crossover transmembrane helices 4 and 5 which brings NBD2 closer to NBD1. The greater rotation of ∼15° of TMH4 and 5 results in a narrower separation of the non-canonical (degenerate) composite ATP binding site (Fig 2B). Based on models built to the dephosphorylated chTS density, the distance between equivalent human NBD1 Walker A residue G461 and NBD2 signature sequence residue S1347 at the non-canonical ATP binding site is calculated to be ∼21 Å compared to the 27 Å reported for zfCFTR (Zhang, Liu et al. 2017). At the canonical composite site, the NBD1 signature sequence S549 is separated from NBD2 Walker A G1247 by ∼16 Å, approximately 2 Å closer than observed for the corresponding residues of dephosphorylated zfCFTR (Fig 2B). Moreover, density corresponding to an adenosine nucleotide is present in both non-catalytic NBDs.

**Figure 3.**
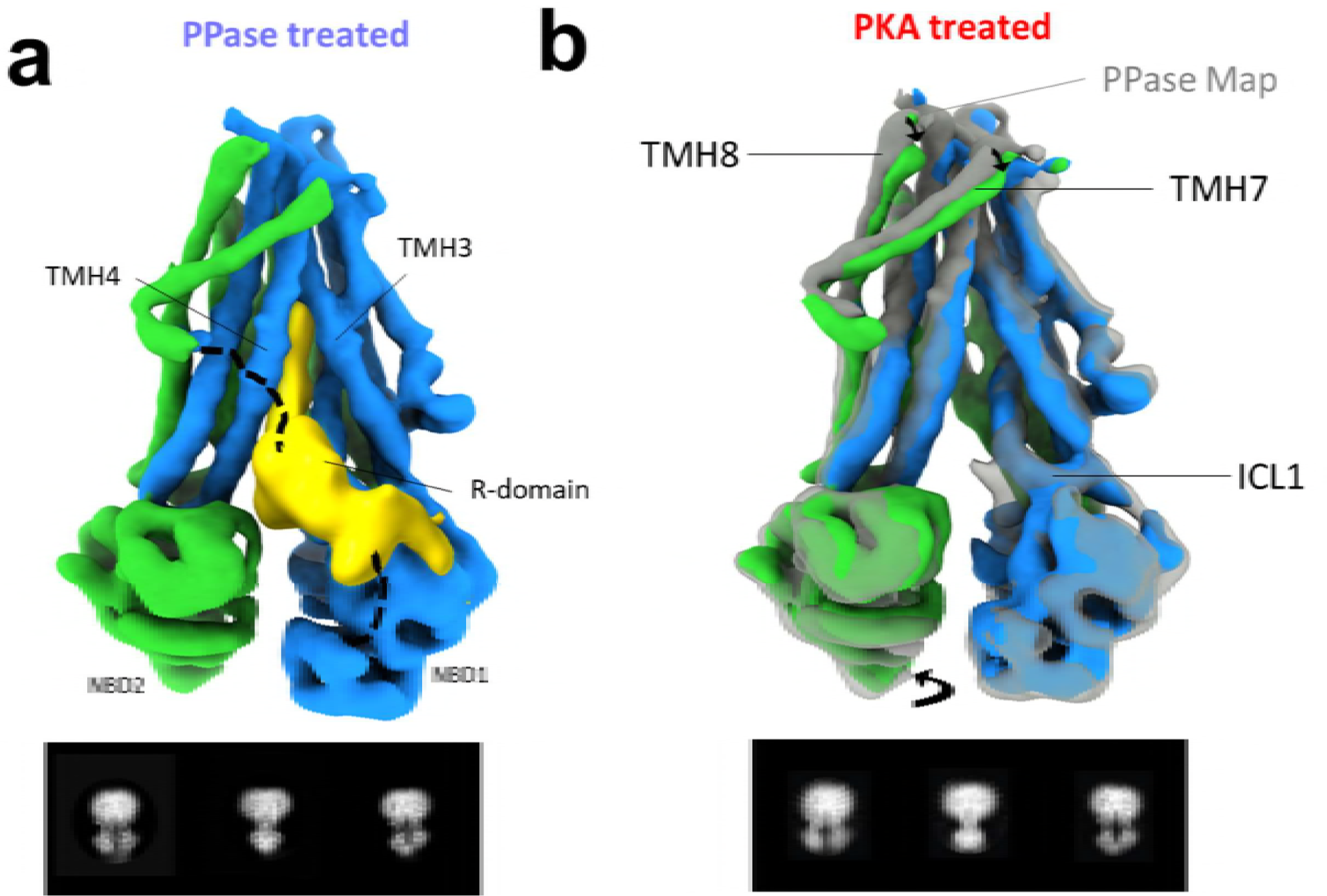
Putative R-domain density engages CFTR in a unique way only in the PPase treated cryo-EM map. **a)** Side views of the unfiltered dephosphorylated chTS cryo-EM map reconstruction with EM density corresponding to the R domain shown in yellow. Note the density extends deep into the intercellular vestibule, as if to act as a “plug”. bottom Representative two-dimensional class averages of dephosphorylated CFTR channels **b)** For comparison side views of the phosphorylated chTS cryo-EM map, colored as in Fig 1B and compared to the translucent gray overlay of the dephosphorylated chTS. Overall, the architecture is similar between the two treatment conditions for chTS. Notably, there is a lack of observable R-domain density after phosphorylation. Small differences are observed in the positions of TMH8 and 7 which appear to have an ∼2 Å rigid body shift towards the cytoplasm (denoted with an arrow). bottom Representative two-dimensional class averages of phosphorylated chCFTR channels.

### Membrane domains

Whereas the conformation of most of the TMD regions are congruent with that in the unphosphorylated, ATP-free zebrafish CFTR, helices 7 and 8 appear to be arranged quite differently. Notably, this positioning of these helices reveals an externally oriented inter-TM opening or vestibule (Fig 2C). This newly observed membrane exposed crevasse has a SiteMap site volume of ∼1400 Å^2^ and is formed by the inner faces of TMHs 5,6,9, and 12.

A distinct difference between our cryo-EM structures and other recently published CFTR cryo-EM structures is that trans-membrane helix 8 (TMH8) does not exist in a kinked conformation (Zhang and Chen 2016; Liu, Zhang et al. 2017; Zhang, Liu et al. 2017) (Fig 2D). The kinked conformation of TMH8 seen by Chen and colleagues occludes the vestibule we observe. Our data places TMH8 in an extended conformation (Fig 2C). For the TMH8 extended conformation to exist, attached extracellular loop 4 (containing the two N-glycosylation sites N894 and N900) and TMH7 would have to rearrange. Indeed, TM7 appears to exist in a novel location as well. Clear density which we interpret to be TMH7 is seen located closer to TMH5 and 4, different from the more canonical location close to TMH9 observed in the CFTR structures of Chen and colleagues and other ABC exporters (Fig 2D). Thus, this TMH7 position is atypical. Consistent with this location is that of putative glycosylation density observed on EL4 at lower threshold density.

### R domain

We observe considerable density that we interpret to be the R-domain, a portion of which is interleaved between the cytoplasmic extensions of TMH3 and TMH4 (Fig 3A). This electron density is too weak to accurately assign secondary structure. Qualitatively, however, there appears to be some interaction of this poorly resolved region with TMH3, the first intracellular loop one (ICL1), and with NBD1 itself, specifically near the ATP binding site and near where the RI would be located. A portion of this putative R-domain density, extends deep within the cytoplasmic core of the transmembrane helical bundle, reaching near human equivalent residue W1145 in TMH12. This arrangement is quite different from that of the R domain density of zfCFTR which interacts primarily with other cytoplasmic domains before a C-terminal segment protrudes between the inner ends of TMHs 9, 10, and 12.

### Phosphorylated chCFTR

Purified PKA treated chCFTR, confirmed to be phosphorylated and activated (Fig 1), was similarly examined by cryo-EM in the presence of 5mM Mg-ATP, however, at lower magnification. From 3388 micrographs, ∼700,000 initial objects were selected, 331,487 particles were retained after “2D filtering” in RELION2 (Scheres 2012). Single-particle three-dimensional analysis of these remaining particles in cryoSPARC (Punjani, Rubinstein et al. 2017) revealed a single class with an overall resolution of 7.62Å (FSC = 0.143). After multiple rounds of 3D classification 57,826 particles could be reconstructed to a 3D map for which the cryoSPARC estimated FSC measurement at 0.143 reached 6.55 Å for the masked protein (Fig 3B, Fig S4).

### Structural Architecture of phosphorylated and comparison with dephosphorlated inactive protein

For the phosphorylated chCFTR reconstruction the transmembrane helices were well resolved. Even though extracellular loop density, was less well resolved than that of the dephosphorylated protein, at lower threshold values the putative EL4-associated-glycosylation density could still be observed. Otherwise the maps were similar except that the density corresponding to the R domain was not observed after PKA treatment. This loss of observable R domain density on phosphorylation is already evident when comparing the 2D class averages of PPase treated and PKA treated (Fig 3) samples. Interestingly, the lack of observable R-domain density between the MSDs in the latter specimen does not result in any major difference in NBD separation, when compared to our dephosphorylated map. A minor reduction in separation is detectible, however, accompanied by a small movement of cross over helices 4 and 5. Thus, it is clear that phosphorylation does not appear to induce stable NBD dimerization in our sample, in contrast to what is seen in the published zebra fish CFTR cryo-EM models (Zhang, Liu et al. 2017).

### Membrane domain variations

Subtle differences between the maps of the dephosphorylated and phosphorylated states is evident in the membrane helices. Comparing the two maps, extra density (presumably corresponding to the R domain) is visible around iCL1 in the PPase map as discussed above (Fig 3B). The very different dispositions of TMHs 7 and 8 from those of those in the structures of Chen and coworkers is retained by chCFTR after phosphorylation. There are, however minor changes in the location of these helices where the phosphorylated models exhibit a pronounced ∼2Å rigid body movement of them away from their locations in the models of the dephosphorylated protein (Fig 3B).

Another notable observation within the membrane domains involves TMHs 12 and 6, known to line opposite sides of the pore. Even at a modest overall 6.55 Å resolution reconstruction, in a better resolved local region, we see clear side chain density in the PKA treated chTS of residue W1149 (1145 human) on TM12, deep within the transmembrane helical bundle. At lower threshold values some density for presumably R353 (R352 human) on TM6 was also present (Fig S5). Densities for these residues in our PPase maps are also observed. The detection of these important residues in the lower resolution PKA treated maps may suggest some degree of structural constraint in this key region. Based on atomic modeling using the map density as a constraint, it becomes obvious that the connection between the intracellular vestibule and the newly observed extracellular vestibule may occur at this location where these two residues in TMH6 and 12 are opposite each other. The implications of these observations are discussed in further detail below.

## Discussion

In this work we focused on CFTR channel protein that was inactive when dephosphorylated and active when treated with protein kinase A to phosphorylate multiple sites on the R domain. We employed chicken CFTR with thermally stabilizing mutations in NBD1 and substitution of the catalytic site histidine in NBD2 (illustrated in Fig S2B) because this construct matured well when stably expressed in mammalian cells, could be purified in a mono-disperse form and was active after reconstitution into proteoliposomes (Fig 1C). Cryo-EM maps of the dephosphorylated and phosphorylated protein, both in the presence of Mg-ATP, showed similarities but also striking differences from those of zebrafish and human CFTR structures (Zhang and Chen 2016; Liu, Zhang et al. 2017; Zhang, Liu et al. 2017). Poorly resolved R-domain densities, a portion of which projected between different cytoplasmic extensions in our maps and those of Chen and colleagues, were undetectable in both cases after phosphorylation by PKA (Fig 3). However our maps showed major differences from those from the Chen group in the relative positioning of the NBDs and the arrangements of the transmembrane helices, particularly TMHs 7 and 8 (Fig 2,4B).

**Figure 4.**
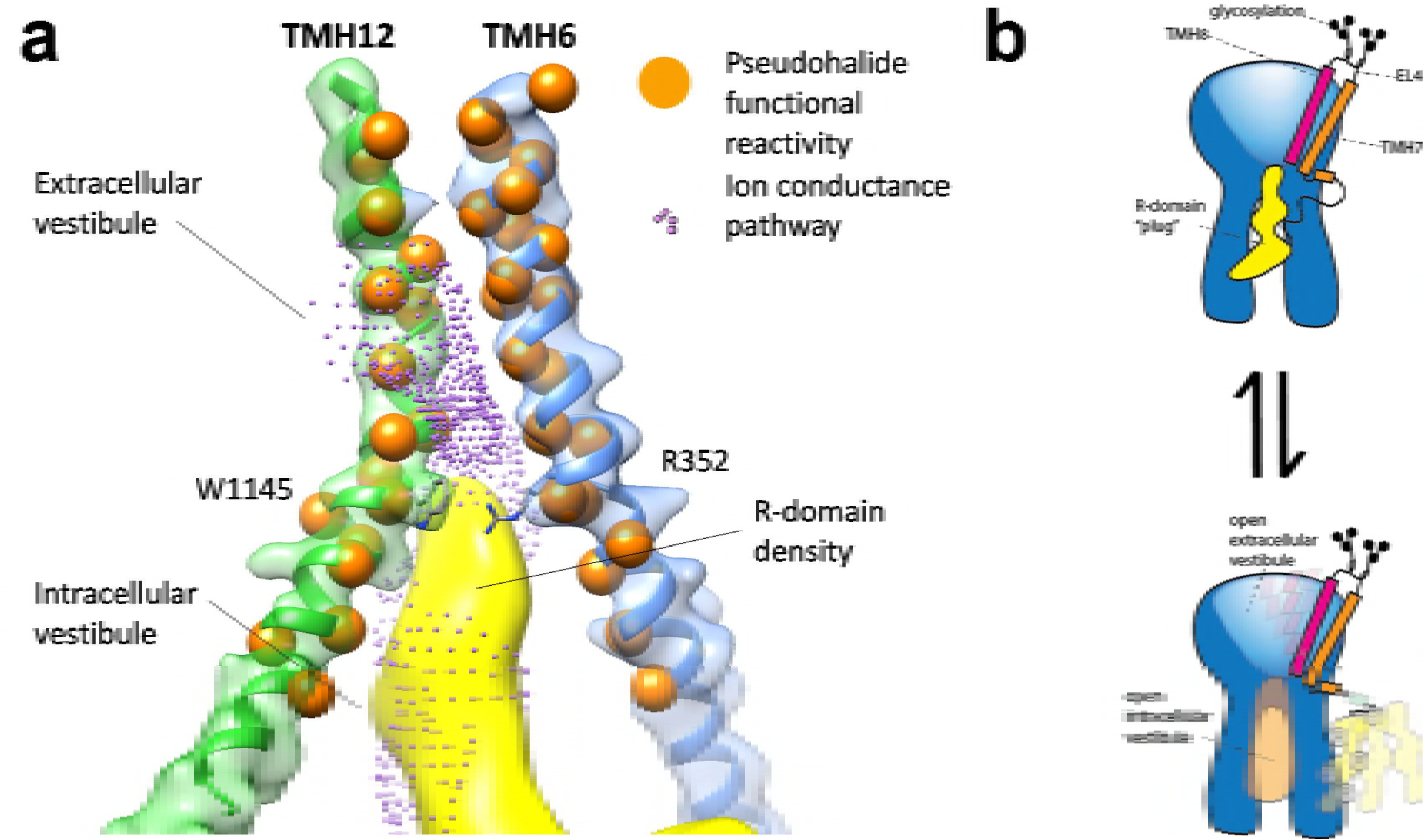
R-domain density passes through the intracellular vestibule and reaches the ion conductance pathway between TMH 12 and 6. **a)** As shown here the R domain (yellow) extends deep into the two halves of CFTR, filling the intracellular vestibule, where it could act as a “plug” limiting Cl ion passage to the newly observed extracellular vestibule. Extracellular permeant pseduohalide cysteine reactive pore lining residues on TMH6 (blue) and TMH12 (green) are shown in orange ((Norimatsu, Ivetac et al. 2012), see below), and a model of the ion conduction pathway calculated using hole2 is shown in plum. Also shown are residues W1145 and R352 for which we observe density for in our map of the phosphorylated active protein. **b)** Cartoon illustrating different structural changes relevant to activation of CFTR channels. CFTR shown in blue has TMH7 (orange) in an atypical ABC protein orientation and the kinked TMH8 represented as series of connected diamonds or an extended conformation strait tube (magenta). **chCFTR** Thermostabilized CFTR has an atypical TMH7 and extended TMH8 conformation that exposes an extracellular vestibule. The connected R-domain (yellow) that enters from between TMH3 and 4 (depicted as entering in front) moves out from between the cytoplasmic extensions of the TMD upon PKA treatment. Thus connecting the large extracellular vestibule (light blue) to the intracellular vestibule (light orange). These movements together may lead to further opening events deep within the vestibule.

### NBD *separation*

Typically, ABC transporters are thought to shift from an inward facing (IWF) conformation with NBDs separated and the TM helices closely associated to an outward facing (OWF) state with NBDs dimerized and rearrangement of TMs thus enabling substrate to be exported across the membrane. Importantly, the cargo is exposed to either only the intracellular environment in the IWF conformation or only the extracellular environment in the OWF conformation. This “alternating access” model of activation has been applied to CFTR activation where the NBDs form a transient “nucleotide-sandwich” state on the pathway to an open channel state (Gadsby, Vergani et al. 2006). Indeed, the phosphorylated zfCFTR structure with a catalytically inactive NBD2 may appear to exist in such a proposed state (Zhang, Liu et al. 2017). However, an open pore through which chloride ions could pass is not apparent in that structure. To function as a channel, a pore would be required to exist perhaps as an intermediate between IWF and OWF conformations with an open connection between the extracellular and intracellular compartments.

The cryo EM maps we have resolved of a high open probability, functional channel, that appears to show extracellular access to the pore, clearly does not have a tight NBD dimer interface. However, an activated state, represented by the observed phosphorylated cryo-EM map present here, may mimic an outward facing state of the more extracellular portions of the TMDs. This spoofing of an outward facing state by TMH rearrangements, specifically, rearrangements of TMH 7 and 8 that reveal an external access to the pore, could arguably be consistent with CFTR co-opting the typical ABC transporter structural framework to mediate regulated passive Cl ion permeation. The smaller ion transport cargo and fast flicking channel activation events would require rapid conformational changes but not necessarily the formation of a fully-realized outward facing conformation.

Regardless, in both zfCFTR and chTS phosphorylated structures, access to the intracellular vestibule is possible either as a portal seen in the “nucleotide sandwich” conformation of zfCFTR or the NBD separated state we observe. Thus, in these two models the state of NBD dimerization would not appear critical for chloride access to the intracellular vestibule. Moreover, previous studies that deleted NBD2, showed PKA dependent but ATP independent gating, suggesting that NBD dimerization is not a requirement for an open channel state (Cui, Aleksandrov et al. 2007; Wang, Bernard et al. 2007; Wang, Wu et al. 2010). Perhaps more importantly is the appearance of an extracellular vestibule that we observe, discussed in further detail below.

### Extracellular vestibule

In the zebrafish and human CFTR structures, transmembrane helix 8 (TMH8) appears in a novel kinked conformation. This kinked region of TMH8 occupies space that for other ABC proteins overlaps with the upper part of a “typical” TMH7. Of the increasing number of ATP-binding cassette (ABC) protein structures determined by crystallography (Locher 2016) and more recently by cryo-electron microscopy (cryo-EM) (Kim, Wu et al. 2015; Frank, Shukla et al. 2016; Oldham, Grigorieff et al. 2016; Oldham, Hite et al. 2016; Zhang and Chen 2016; Fitzpatrick, Llabres et al. 2017; Johnson and Chen 2017; Liu, Zhang et al. 2017; Martin, Yoshioka et al. 2017; Mi, Li et al. 2017; Qian, Zhao et al. 2017; Taylor, Manolaridis et al. 2017; Zhang, Liu et al. 2017), none show the unique conformation that we observe in our cryo-EM 3D reconstructions presented here. Notably, TMH8 appears in a more conventional extended conformation and not a kinked conformation as seen in the recent cryo-EM maps, reported by Chen and colleagues. When aligned to MSD2 we see that the different conformation of TMH8 in our structure exists in an extended state, similar to the TMH8 of an OWF of SAV1866 (Dawson and Locher 2006). Strikingly, we observe density for TMH7 in a conformation previously unseen for this helix in ABC proteins, as our TMH7 density is located near TMH4 in our reconstructions (Fig 2C,D). Additionally, the non-canonical rearrangement of TMH7 observed in our structures could support the extended conformation of TMH8. In this way CFTR can mimic an outward facing state that is observed for SAV1866, with only a rearrangement of TMH7 and 8, leading to formation of a large vestibule observed between TMH5,6,9 and 12 (Fig 2C). These observations prompt consideration of possible novel structural mechanisms of CFTR activation discussed in more detail below.

### Plausible chloride permeation route

The extended conformation of TMH8 we observed instead of the kinked conformation seen by Chen and colleagues (Zhang and Chen 2016; Liu, Zhang et al. 2017; Zhang, Liu et al. 2017) reveals an extracellular vestibule and hints at a possible extracellular chloride access route.

Indeed, TMH8 has been previously shown to be important for CFTR function and biosynthesis. Residue, D924 in TMH8 has been proposed to form a functionally important salt bridge with R347 on TMH6 (Cotten and Welsh 1999; Cui, Freeman et al. 2013). More recently this salt bridge is reported to be observed in the near-atomic cryo-EM map of ATP-free dephosphorylated zfCFTR, although there is not density for the 924 side chain (Kumar and Nussinov 2002; Zhang and Chen 2016). Cystic Fibrosis causing mutations at residue 347 (R347C/H/L/P) suggest the functional importance of this constraint. In agreement, the D924R mutation in TMH8 leads to abnormal gating (erratic flickery) presumably through disruption of this hypothetical salt bridge (Cotten and Welsh 1999). However, Cui and coworkers concluded that the salt bridge made at most a weak contribution to the fully open channel state (Cui, Freeman et al. 2013). The extension of TMH8 out away from the previously observed kinked position would appear to preclude formation of the proposed R347-D924 salt bridge. However, in our structures and those of zfCFTR the upper portion of TMH8 is of generally weaker density than the majority of the TMHs, suggesting this region of TMH8 is a dynamic and thus may populate other alternative states in the cryo-EM ensemble.

Although it is not obvious why the positioning of TMHs 7 and 8 in our structures is different from that in the structures of Chen and colleagues, it is worth recalling that the co-translational incorporation of these two helices into the membrane during biosynthesis occurs by an atypical mechanism involving an unusually high degree of flexibility and cooperativity between them (Carveth, Buck et al. 2002; Kim and Skach 2012). While TMH 7 has conventional signal sequence activity, TMH8 is only to able exhibit stop transfer activity when paired with TMH7. TMH8 appears mobile within the membrane during incorporation as evidenced by its extending to variable degrees into the ER lumen as reflected by accessibility to oligosaccharide transferase of asparagine residues in EL4 at different distances from the membrane (Carveth, Buck et al. 2002). During post translational processing the two core N-linked oligosaccharide chains in EL4 are trimmed and extended to form large multi-antennary complex chains. Since increasing the hydrophobicity of TMH8 is known to “pull” the helix deeper into the membrane one might speculate that addition of the large polar carbohydrate chains to EL4 might “pull” the helix in the opposite direction. In this regard it is interesting to note that our CFTR constructs used here have the large complex oligosaccharides whereas those of Chen and coworkers have only the short core chains because of the different expression systems used.

Overall we observe that membrane spanning domain 2 (MSD) appears to have a lower local resolution than MSD1 (Fig S3), as is also observed in zfCFTR cryo-EM reconstructions. In addition to variability in the TMH7-EL4-TMH8 unit arising during biogenesis we specifically observed weaker EM densities in this unit as was also observed for TMH8 in the ATP bound phosphorylated zebrafish CFTR cryo-EM structure. These findings are consistent with this being a highly mobile region that may be critical in channel function.

In the lower resolution phosphorylated map, we cannot place many of the side chains with certainty, and therefore we cannot definitely comment on the fine details of the phosphorylated state pore. However, close examination of the higher resolution dephosphorylated map reveals a pore at the connection point between the intracellular and extracellular vestibule.

We speculate that the previously unseen large extracellular vestibule in our structure may provide extracellular access to the pore. Of the side chains we can observe in both maps, two that appear deep in the transmembrane helical bundle at the bottom of this vestibule correspond to human residues W1145 and R352 (Fig 4 and Fig S5). Interestingly, sulfhydryl modification of W1145C is inhibitory to channel function and crosslinking through use of a double cys mutant W1145C/R352C leads to a channel that is also nonfunctional in the absence of reducing agent, yet is functional in the presence of reducing agent (Das, Aleksandrov et al. 2017). Notably, these residues lie around the narrowest part of the putative pore, in atomic models based on the cryo-EM map density we report here, and thus may play a fundamental role in normal channel gating. Interestingly, we observe a π-helix bulge between human residues R347 and R352, in TMH6 in our dephosphorylated CFTR map. While we cannot confirm a π-helix bulge at our lower resolution phosphorylated cryo-EM map, we do observe low threshold density of residue R352 near this critical region. The energetically unfavorable nature of a π-helix bulge and the proximity of key residues, suggest that this region may be a structural trigger point for propagation of gating conformational transitions.

Interestingly, TMH6 and/or 12 line the substrate translocation pathway of ABC exporters. Moreover, these key TMHs are both seen to become less alpha helical in many ABC protein structures. We also observe putative lipid densities on the outer face of one of these two TMHs (Fig S6). In CFTR TMH6 and 12 have been the subject of intense investigation by cysteine-scanning mutagenesis (reviewed in (El Hiani and Linsdell 2014; Wang, El Hiani et al. 2014)). While extracellular MTS reagents seem to penetrate poorly, studies from Dawson and colleagues using permeant, pseudohalide anions (eg Ag(CN)_2_) have shown reactivity deeper than TMH6’s R352 and TMH12’s W1145 (reviewed in (Norimatsu, Ivetac et al. 2012); Fig 4A, orange balls), thus lending support to the idea that the newly observed external vestibule we observe (formed by TMH5,6,9,&12) may enable permeation to the level of this residue pair. Based on atomic models built from our density, it appears likely the connection between the intracellular and extracellular vestibules may occur between TMH6 and 12 around the level of residues R352 and W1145 (Fig 4A). This proposal is consistent with chemical modifications of cysteine replacements of residues that line these two critical TMHs having modulatory effects on CFTR function (Bai, Li et al. 2010; El Hiani and Linsdell 2010; Bai, Li et al. 2011; Norimatsu, Ivetac et al. 2012).

### R domain

CFTR is a unique ABC family member by virtue of its function as an anion channel, rather than an active transporter. Moreover, unlike the other members of this family, CFTR contains a unique large regulatory (R) domain that must become phosphorylated for the channel to be activated. Notably, at least a part of the R domain’s critical role in channel activation has been proposed to be independent of ATP-induced NBD dimerization (Wang, Wu et al. 2010). In fact, deletion of this key domain (Rich, Gregory et al. 1991) or a portion of it (residues 817–838) (Xie, Adams et al. 2002) completely eliminates the PKA dependence of channel activity.

In our dephosphorylated map the R domain density that extends into the intracellular vestibule penetrates right to its connection with the extracellular vestibule (Fig 4A, yellow). Thus one could envision the R domain density acting as a “plug”, blocking chloride permeation through this constriction. TMH7 is located directly adjacent to and is linked to the R-domain. In our active state maps, TMH7 and 8 move by about 2Å relative to their position in our inactive state map, suggesting that release of the R domain from between the two halves of CFTR could allow for further adjustments of TMH7 and 8 in addition to the subtle NBD contraction. Thus, it is tempting to speculate based on our cryo-EM structures that phosphorylation of the R-domain allows a portion of it to transition from acting as a blocker of permeation to an unstructured region that is no longer observed and thus unable to block permeation (Fig 4B).

In consideration of this proposed role of the R domain, it is important to emphasize that our proposed R domain density is different from that of Chen and colleagues, that extends out between TMH9, 10, and 12. Instead we see this density take a different route, where it penetrates deep into the transmembrane helical bundle between the TMDs entering on the opposite side of the protein between TMH 3 and 4 (Fig 3A). The differential entry of the R-domain is due to the alterative positions we observe for TMH7.

### TM7 dynamics and linkage of alternative structures

An illustration of the major variances between the phosphorylated and dephosphorylated chCFTR is shown in Figure 4B. When considering only architecturally how these alternative structural states could arise, in relation to the zfCFTR structures, the linking of these states could only seem to occur after the R domain is unconstrained. This is due to the different arrangement of TMH7 between zfCFTR and chCFTR and differential R domain engagements. Thus, the link between these alternative structures could occur most readily between the two alternative PKA treated states of zfCFTR and chCFTR. In this case, simplistically, the R domain would release from between the two halves of CFTR leaving its connected TMH7 free to explore alternative states. Subsequent dephosphorylation after TMH7 rearrangements could facilitate the differential re-engagement of the R domain, which is observed between the quiescent ATP free and ATP containing, dephosphorylated inactive states.

### Conclusion

Here we present a unique CFTR structural state that is not only new to CFTR but also to the ABC protein family. While we cannot conclusively state that the phosphorylated cryo-EM map we observe illustrate an open pore state due to limitations in the current resolution, ongoing and future studies seek to address these atomic details. Moreover, the differences we observe in our CFTR structures compared to those from Chen and colleagues underscores the importance of continued structural biological efforts to resolve different CFTR structural states to gain a greater appreciation of the rugged and dynamic landscape of CFTR.

## Materials and Methods

### Expression and purification of ABCC7

Expression of Chicken ΔRI/1404S/1441X/His construct was performed as previously described (Aleksandrov, Jensen et al. 2015). In brief, suspension adapted BHK cells stably expressing this construct were grown in SFM II 293s suspension medium (Invitrogen) supplemented with 2% FBS, 2x glutamine, and 250 μM methotrexate in a 2 L square PETG plastic medium bottles (Fisher) at 37°C at 120 rpm.

All subsequent steps were performed on ice or 4°C unless otherwise noted. Cells were pelleted in PBS, lysed in hypotonic buffer (50 mM HEPES pH 7.4, 1 mM EDTA) supplemented with protease inhibitor cocktail [PIC: leupeptin (Research Products International, RPI) 9.0 μg/ml, aprotinin, (RPI) 12.0 ?μg/ml, Pefabloc (RPI) 450 μg/ml, benzamidine (Fisher), 100 μg/ml, E64 (AG Scientific) 30 μg/mL], with a glass mortar and pessle (Kontes Glass). Lysed cells were osmotically balanced by the addition of 250 mM final sucrose, and cell debris was removed by centrifugation at 4200 x g. The supernatant was then pelted at 125,000 x g for 30 min and resuspended in 30 mM Tris–HCl, pH 7.4; 0.5 M NaCl; 0.1 M arginine; 20% glycerol, 5mM ATP-Mg with PIC. A pre-clearing step was performed by incubating resuspended membranes with 0.01% DMNG for 15 min and further centrifugation at 125,000 x g for 30 min. Cleared membrane pellet was then resuspended in sodium phosphate buffer [50 mM Na-PO4, pH 7.6, 0.5M NaCl, 0.1 M arginine, 15% glycerol, PIC, 5 mM ATP-Mg] and solubilized with 1% DMNG for 30 min. Solubilized extract was cleared by centrifugation at 125,000 x g for 30 min and applied to Co-Talon resin (Clontec) for 3-4 hours. Resin was washed extensively with sodium phosphate buffer containing 25-50mM imidazole with 0.1% DMNG. Resin was then exchanged into wash buffer [40 mM Tris–HCl, pH 7.4; 0.15 M NaCl; 0.1 M arginine; 10% glycerol, 5 mM ATP-Mg] supplemented with 0.01% DMNG. PKA (VWR) or lambda (Fisher) and alkaline phosphatase (NEB) treatment was performed on the beads at ∼100 units/mL supplemented with 1mM manganese where appropriate for >16hrs at 4°C. Treated CFTR was then detergent exchanged into 0.06%/3mM Digitonin (Sigma) / Fluorinated Fos-Choline-8 (Anatrace) in wash buffer. Importantly, 5-10% Digitonin stock was prepared >16 hours prior, and filtered the next day. The critical micelle concentration of the filtered Digitonin stock was empirically determined using a DPH fluorescent assay (Chattopadhyay and London 1984) and typically “0.06%” represented between 2-3 times the CMC. CFTR was eluted in elution buffer with 200 mM imidazole, concentrated to ∼7 mg/mL using 150 kDa cut off filters (Apollo 7mL, OrbitalBioSciences). Finally, concentrated CFTR samples were buffer exchanged into EM sample buffer [40 mM Tris–HCl, pH 7.4; 0.15 M NaCl; 5mM ATP-Mg; 0.06% Digitonin; 3 mM FFC8].

### Liposome preparation for single channel recordings

Preparation of CFTR containing proteoliposomes was performed as previously described (Aleksandrov, Jensen et al. 2015). In brief, Large Uni Lameller Vesicles (LUVS) comprised of DOPE, DOPC, DOPS, Cholesterol (48:22:12:18) respectively were destabilized with Triton X-100 prior to addition of purified PKA treated CFTR at a 1:50 ratio of protein to lipids and detergent removal by Bio-Beads SM2.

### Single channel analysis

Single channel recordings were performed as previously described (Aleksandrov, Jensen et al. 2015). In brief, planar lipid bilayers (3:1 mix of POPE and POPS) were spontaneously fused with CFTR ion channels contained in proteoliposomes in the presence of 5 mM ATP-Mg, 50 nM PKA to the cis chamber. Symmetrical salt solution was maintained using a buffer of 300 mM Tris-HCL, pH 7.2, 3 mM MgCl_2_ and 1mM EGTA and voltage-clamp recordings were recorded. All-points histograms (pCLAMP 9.2, Axon Instruments) were fit with Origin 4.1 software (OriginLab Corp., Northampton, MA, USA) and probability of the single channel being open (P_o_) determined as from the ratio of the peak integrals.

### Phosphorylated Cryo-EM grid preparation, data acquisition and Image processing

Ice cold purified Chicken ΔRI/1404S/1441X/His construct (chTS) CFTR at 3-4 mg/mL in EM sample buffer was added (5 uL) to glow discharged grids (10 mA, glow time 1-2 min) using a PELCO easiGlow Discharge Cleaning System. Quantifoil R1.2/1.3 400 mesh Cu holey carbon grids were blotted (∼3.5 seconds) with Vitrobot Mark IV (FEI) and plunge frozen in liquid ethane

Images were collected using a 300keV TitanKrios (FEI) with a Gatan K2 Summit direct electron detector in SuperRes mode at a physical pixel size of 1.71 Å. 4283 micrograph recorded images were automatically collected using Serial EM using GIF (Gatan Imagining Filter) with a slit width of 20 ev. Data were collected at a dose rate of ∼5 e/pix/sec as recorded from counting mode. Images were recorded for 25 seconds in 100 subframes to give a total dose of ∼42 electron per Å^2^.

Movie frames devoid of gain reference were compressed >5 fold using Zstandard (Yan Collet, Facebook) to assist in data transfer for off-site data processing. Super-res movie frames were gain reference subtracted and aligned using MotionCorr (7x7 patch) to correct for particle drift motion (Zheng, Palovcak et al. 2017). Aligned micrographs were Fourier binned by 2 using Imod. The contract transfer function (CTF) of each micrograph was estimated using gCTF (Zhang 2016). Manual inspection of the micrographs removed about 25% (only carbon, thick-ice, poorly motion corrected, etc.). The remaining micrographs had a calculated mean defocus of 3.0 μm (SD = 1.0?μm). Particles were automatically selected from the aligned micrographs using a difference of Gaussians (DoG) particle picker (Voss, Yoshioka et al. 2009), accelerated with GNU parallels (Tange 2011) and also re-picked using a Gaussian filtered blob in PARTICLE (J.Z. Chen). The average of overlapping picked particles from the two sets were merged along with the re-merging of a subset of ∼200k top particles selected from four rounds of single elimination Relion2.0 2D and 3D classification yielding about 700k particles from 3388 micrographs, using a general purpose particle merging utility (J.Z. Chen). After 1 round of double elimination two-dimensional classification in Relion2.0 and selection, a final data set of 331,487 particles were obtained. All subsequent three-dimensional classification and refinement steps were performed within cryoSPARC (Punjani, Rubinstein et al. 2017). Five rounds of multi-reference refinement with two to four classes resolved 57,826 particles that produced a map with a resolution of 6.6?Å (by FSC using the 0.143?Å cut-off criterion) (Rosenthal and Henderson 2003) after post-processing including soft masking and B-factor sharpening (B-factor of −554.2?Å^2^) in cryoSPARC.

### Dephosphorylated CFTR Cryo-EM grid preparation, data acquisition and Image processing

Ice cold purified PPase treated Chicken ΔRI/1404S/1441X/His construct (chTS) CFTR at 7-8 mg/mL in D_2_O containing EM sample buffer was added (5 uL) to glow discharged (15 mA, glow time 90 sec) using a PELCO easiGlow Discharge Cleaning System. Quantifoil R1.2/1.3 400 mesh Au holey carbon grids, blotted (4-6 seconds) with Vitrobot Mark IV (FEI), and plunge frozen in liquid ethane

The PPase data set consisting of 783 images collected as described above, except the images were recorded in SuperRes mode at a physical pixel size of 1.37 Å for 20 or 10 seconds at 5 e/pix/sec or 7.5 e/ pix/sec at 10 frames a second to give a dose of ∼55 or 40 electrons per Å^2^, respectively. The energy filter slit width was also set to 10 ev. The 754 remaining micrographs after manual curation had a calculated mean defocus of 2.3 μm (SD = 0.7?μm). 221,536 particles were automatically selected using template picking and refined to 165,541 after 1 round of single elimination two-dimension classification in RELION 2.0. Per particle defocus, 2D filtered particles were then subjected to at least five rounds of multi-reference refinement with two to four classes to resolve 30,219 particles that produced a map with a cryoSPARC calculated resolution of 4.79?Å (soft whole particle masking). Post-processing included B-factor sharpening (B-factor of - 168.2?Å^2^) in cryoSPARC. Of note, application of the K2 MTF prior to B-factor sharpening resulted in a Guiner estimated B-factor of -98.12 Å^2^. As the Guinier estimation performed in cryoSPARC alone did not resolve higher-resolution features, the resulting map was then spectrally filtered from 10 to 2.6 Å (Grigorieff 2018), and then filtered by local resolution (using blocfilt, kernel size 11) (Heymann and Belnap 2007)) to generate the penultimate post-processed sharpened map (Fig S3E).

Next, the 30,219 particles were re-extracted from super resolution images (0.68 Å/pix). An additional round of 3D classification retained ∼95% of the super-res particles. The spectrally flattened local resolution filtered penultimate map (1.37 Å/pix) was used as a template with refine_res_init set to 5 Å for subsequent refinement in cryoSPARC. The resulting map gold standard FSC resolution of for the masked protein was 4.3 Å (Fig S3).

### Model building

A manual rebuild of TMH8 and rigid body docking TMH7 into the EM map density was performed using COOT (Emsley, Lohkamp et al. 2010) and used as a down weighted (0.7) template along with PDB ID code 5UAR as the primary template for input into RosettaCM to generate 100 models. Specifically, the TM8-EL4-TM7 region (from residues 848 to 953) was analyzed in detail and the highest overall aggregate scoring (EMringer (Barad, Echols et al. 2015), MolProbity (Williams, Headd et al. 2018), Rosetta Score and fit to density (elec_dens_fast) model was selected. The TM8-EL4-TM7 region model was then refined using an iterative local rebuilding and atom refinement as previously described (Wang, Song et al. 2016) with 2 additional rounds of LocalRelax. 500 models were generated and the best normalized aggregate score for the TM8-EL4-TM7 region model was selected and used as an additional template for additional rosettaCM model generation outlined below.

Next, PDB ID codes 5UAR & 5W81 were used as a starting template models (Zhang and Chen 2016; Zhang, Liu et al. 2017). RosettaCM was then used to generate ∼500 chCFTR PPase treated models using density as a constraint (Song, DiMaio et al. 2013) and updated sidechain rescaling factors (Wang, Song et al. 2016). Missing loop regions (i.e. EL4, R-domain, TMH12 linker to NBD2) were removed for analysis and one of the top 5 models with highest overall combined normalized aggregate score (EMringer (Barad, Echols et al. 2015), MolProbity (Williams, Headd et al. 2018), Rosetta Score and fit to density (elec_dens_fast)) was selected. This model was then used as a seed for a second round rosettaCM from 500 models the top 50 highest scoring integrated FSC from 10-4Å (iFSC) was selected, and then the model with the best MolProbity score was selected as the final model.

The resulting selected model was used to generate 500 models via iterative local rebuilding and atom refinement (Wang, Song et al. 2016). Then the top 50 models with the best iFSC were selected and the top 10 with the best MolProbity score were then used to plot RMSD at Cα to illustrate model uncertainty (Fig S7).

The PKA treated model was constructed by taking the PPase chCFTR model and fitting it into the PKA treated map density using iModfit (Lopez-Blanco and Chacon 2013). The final PKA model side chains were made “stubby” by deleting atoms past the β carbon. Residues 1149 and 353 were retained in the final model as they had reasonably well-defined side-chain densities past the β carbon (Fig S5).

## Data availability

Atomic coordinates of the models of chCFTR fit to the respective cryo-EM maps are available under accessions WWW and XXX, for phosphorylated and dephosphorylated, respectively. The three-dimensional cryo-EM density maps (half maps and sharpened) have been deposited in Electron Microscopy Data Bank under accessions EMD-YYYY (phosphorylated) and EMD-ZZZZ (dephosphorylated). Binned Particle stacks and meta data can be found at https://XXX/XXX.

## Acknowledgments

We thank Drs. R. Ford, S. Shyng, S. Mansoor, and G. Martin for discussions and/or critical reading of the manuscript. We are grateful to C. Yoshioka, C. López, the Multiscale Microscopy Core (MMC), Oregon Health & Science University (OHSU)-FEI Living Lab and the OHSU Center for Spatial Systems Biomedicine (OCSSB Multiscale Microscopy Core (OHSU) for their outstanding microscopy support. This work was funded, in part, by CFF fellowship (to J.F.F.), grants RIORDA07XXO and DK051870 (to J.R.R.), and CFF pilot grant (to J.Z.C). The Titan X Pascal used for this research was kindly donated to J.F.F. by the NVIDIA Corporation.

## References

Aleksandrov, A. A., P. Kota, et al. (2012). “Allosteric modulation balances thermodynamic stability and restores function of DeltaF508 CFTR.” J Mol Biol, 419(1-2): 41–60.

Aleksandrov, L. A., T. J. Jensen, et al. (2015). “Thermal stability of purified and reconstituted CFTR in a locked open channel conformation”. Protein Expr Purif 116: 159–166.

Bai, Y., M. Li, et al. (2010). “Dual roles of the sixth transmembrane segment of the CFTR chloride channel in gating and permeation.” J Gen Physiol 136(3): 293–309.

Bai, Y., M. Li, et al. (2011). “Structural basis for the channel function of a degraded ABC transporter,CFTR (ABCC7).” J Gen Physiol 138(5): 495–507.

Barad, B. A., N. Echols, et al. (2015). “EMRinger: side chain-directed model and map validation for 3D cryo-electron microscopy.” Nat Methods 12(10): 943–946.

Carveth, K., T. Buck, et al. (2002). “Cooperativity and flexibility of cystic fibrosis transmembrane conductance regulator transmembrane segments participate in membrane localization of a charged residue.” J Biol Chem 277(42): 39507–39514.

Chattopadhyay, A. and E. London (1984). “Fluorimetric determination of critical micelle concentration avoiding interference from detergent charge.” Anal Biochem 139(2): 408–412.

Cotten, J. F. and M. J. Welsh (1999). “Cystic fibrosis-associated mutations at arginine 347 alter the pore architecture of CFTR. Evidence for disruption of a salt bridge.” J Biol Chem 274(9): 5429–5435.

Cui, G., C. S. Freeman, et al. (2013). “Two salt bridges differentially contribute to the maintenance of cystic fibrosis transmembrane conductance regulator (CFTR) channel function.” J Biol Chem 288(28): 20758–20767.

Cui, L., L. Aleksandrov, et al. (2007). “Domain interdependence in the biosynthetic assembly of CFTR.” J Mol Biol 365(4): 981–994.

Dalton, J., O. Kalid, et al. (2012). “New model of cystic fibrosis transmembrane conductance regulator proposes active channel-like conformation.” J Chem Inf Model 52(7): 1842–1853.

Das, J., A. A. Aleksandrov, et al. (2017). “Transmembrane helical interactions in the CFTR channel pore.” PLoS Comput Biol 13(6): e1005594.

Dawson, R. J. and K. P. Locher (2006). “Structure of a bacterial multidrug ABC transporter.” Nature 443(7108): 180–185.

El Hiani, Y. and P. Linsdell (2010). “Changes in accessibility of cytoplasmic substances to the pore associated with activation of the cystic fibrosis transmembrane conductance regulator chloride channel.” J Biol Chem 285(42): 32126–32140.

El Hiani, Y. and P. Linsdell (2014). “Conformational changes opening and closing the CFTR chloride channel: insights from cysteine scanning mutagenesis.” Biochem Cell Biol 92(6): 481–488.

Emsley, P., B. Lohkamp, et al. (2010). “Features and development of Coot.” Acta Crystallogr D Biol Crystallogr 66(Pt 4): 486–501.

Fitzpatrick, A. W. P., S. Llabres, et al. (2017). “Structure of the MacAB-TolC ABC-type tripartite multidrug efflux pump.” Nat Microbiol 2: 17070.

Frank, G. A., S. Shukla, et al. (2016). “Cryo-EM Analysis of the Conformational Landscape of Human P-glycoprotein (ABCB1) During its Catalytic Cycle.” Mol Pharmacol 90(1): 35–41.

Gadsby, D. C., P. Vergani, et al. (2006). “The ABC protein turned chloride channel whose failure causes cystic fibrosis.” Nature 440(7083): 477–483.

Grigorieff, N. G. T. Rhou, A. (2018). “cisTEM: User-friendly software for single-particle image processing.” bioRxiv doi:10.1101/257618.

Heymann, J. B. and D. M. Belnap (2007). “Bsoft: image processing and molecular modeling for electron microscopy.” J Struct Biol 157(1): 3–18.

Johnson, Z. L. and J. Chen (2017). “Structural Basis of Substrate Recognition by the Multidrug Resistance Protein MRP1.” Cell 168(6): 1075–1085 e1079.

Kim, J., S. Wu, et al. (2015). “Subnanometre-resolution electron cryomicroscopy structure of a heterodimeric ABC exporter.” Nature 517(7534): 396–400.

Kim, S. J. and W. R. Skach (2012). “Mechanisms of CFTR Folding at the Endoplasmic Reticulum.” Front Pharmacol 3: 201.

Kumar, S. and R. Nussinov (2002). “Close-range electrostatic interactions in proteins.” Chembiochem 3(7): 604–617.

Liu, F., Z. Zhang, et al. (2017). “Molecular Structure of the Human CFTR Ion Channel.” Cell 169(1): 85–95 e88.

Locher, K. P. (2016). “Mechanistic diversity in ATP-binding cassette (ABC) transporters.” Nat Struct Mol Biol 23(6): 487–493.

Lopez-Blanco, J. R. and P. Chacon (2013). “iMODFIT: efficient and robust flexible fitting based on vibrational analysis in internal coordinates.” J Struct Biol 184(2): 261–270.

Martin, G. M., C. Yoshioka, et al. (2017). “Cryo-EM structure of the ATP-sensitive potassium channel illuminates mechanisms of assembly and gating.” Elife 6.

Mi, W., Y. Li, et al. (2017). “Structural basis of MsbA-mediated lipopolysaccharide transport.” Nature 549(7671): 233–237.

Mornon, J. P., B. Hoffmann, et al. (2015). “Full-open and closed CFTR channels, with lateral tunnels from the cytoplasm and an alternative position of the F508 region, as revealed by molecular dynamics.” Cell Mol Life Sci 72(7): 1377–1403.

Norimatsu, Y., A. Ivetac, et al. (2012). “Cystic fibrosis transmembrane conductance regulator: a molecular model defines the architecture of the anion conduction path and locates a “bottleneck” in the pore.” Biochemistry 51(11): 2199–2212.

Oldham, M. L., N. Grigorieff, et al. (2016). “Structure of the transporter associated with antigen processing trapped by herpes simplex virus.” Elife 5.

Oldham, M. L., R. K. Hite, et al. (2016). “A mechanism of viral immune evasion revealed by cryo-EM analysis of the TAP transporter.” Nature 529(7587): 537–540.

Punjani, A., J. L. Rubinstein, et al. (2017). “cryoSPARC: algorithms for rapid unsupervised cryo-EM structure determination.” Nat Methods 14(3): 290–296.

Qian, H., X. Zhao, et al. (2017). “Structure of the Human Lipid Exporter ABCA1.” Cell 169(7): 1228–1239 e1210.

Rahman, K. S., G. Cui, et al. (2013). “Modeling the conformational changes underlying channel opening in CFTR.” PLoS ONE 8(9): e74574.

Ren, H. Y., D. E. Grove, et al. (2013). “VX-809 corrects folding defects in cystic fibrosis transmembrane conductance regulator protein through action on membrane-spanning domain 1.” Mol Biol Cell 24(19): 3016–3024.

Rich, D. P., R. J. Gregory, et al. (1991). “Effect of deleting the R domain on CFTR-generated chloride channels.” Science 253(5016): 205–207.

Rosenthal, P. B. and R. Henderson (2003). “Optimal determination of particle orientation, absolute hand, and contrast loss in single-particle electron cryomicroscopy.” J Mol Biol 333(4): 721–745.

Scheres, S. H. (2012). “RELION: implementation of a Bayesian approach to cryo-EM structure determination.” J Struct Biol 180(3): 519–530.

Song, Y., F. DiMaio, et al. (2013). “High-resolution comparative modeling with RosettaCM.” Structure 21(10): 1735–1742.

Tange, O. (2011). “GNU Parallel - The Command-Line Power Too.” The USENIX Magazine 36(1): 42–47.

Taylor, N. M. I., I. Manolaridis, et al. (2017). “Structure of the human multidrug transporter ABCG2.” Nature 546(7659): 504–509.

Vergani, P., A. C. Nairn, et al. (2003). “On the mechanism of MgATP-dependent gating of CFTR Cl-channels.” J Gen Physiol 121(1): 17–36.

Voss, N. R., C. K. Yoshioka, et al. (2009). “DoG Picker and TiltPicker: software tools to facilitate particle selection in single particle electron microscopy.” J Struct Biol 166(2): 205–213.

Wang, R. Y., Y. Song, et al. (2016). “Automated structure refinement of macromolecular assemblies from cryo-EM maps using Rosetta.” Elife 5.

Wang, W., K. Bernard, et al. (2007). “Curcumin opens cystic fibrosis transmembrane conductance regulator channels by a novel mechanism that requires neither ATP binding nor dimerization of the nucleotide-binding domains.” J Biol Chem 282(7): 4533–4544.

Wang, W., Y. El Hiani, et al. (2014). “Relative contribution of different transmembrane segments to the CFTR chloride channel pore.” Pflugers Arch 466(3): 477–490.

Wang, W., J. Wu, et al. (2010). “ATP-independent CFTR channel gating and allosteric modulation by phosphorylation.” Proc Natl Acad Sci U S A 107(8): 3888–3893.

Williams, C. J., J. J. Headd, et al. (2018). “MolProbity: More and better reference data for improved all-atom structure validation.” Protein Sci 27(1): 293–315.

Xie, J., L. M. Adams, et al. (2002). “A short segment of the R domain of cystic fibrosis transmembrane conductance regulator contains channel stimulatory and inhibitory activities that are separable by sequence modification.” J Biol Chem 277(25): 23019–23027.

Yang, Z., E. Hildebrandt, et al. (2018). “Structural stability of purified human CFTR is systematically improved by mutations in nucleotide binding domain 1.” Biochimica et Biophysica Acta (BBA) - Biomembranes 1860(5): 1193–1204.

Zhang, K. (2016). “Gctf: Real-time CTF determination and correction.” J Struct Biol 193(1): 1–12.

Zhang, Z. and J. Chen (2016). “Atomic Structure of the Cystic Fibrosis Transmembrane Conductance Regulator.” Cell 167(6): 1586–1597 e1589.

Zhang, Z., F. Liu, et al. (2017). “Conformational Changes of CFTR upon Phosphorylation and ATP Binding.” Cell 170(3): 483–491 e488.

Zheng, S. Q., E. Palovcak, et al. (2017). “MotionCor2: anisotropic correction of beam-induced motion for improved cryo-electron microscopy.” Nat Methods 14(4): 331–332.

